# Addition of insoluble fiber to isolation media allows for increased metabolite diversity of lab-cultivable microbes derived from zebrafish gut samples

**DOI:** 10.1101/854109

**Authors:** Alanna R. Condren, Maria S Costa, Natalia Rivera Sanchez, Sindhu Konkapaka, Kristin L Gallik, Ankur Saxena, Brian T Murphy, Laura M Sanchez

**Affiliations:** Department of Pharmaceutical Sciences, University of Illinois at Chicago, 833 S Wood St Chicago, IL 60612; Faculty of Pharmaceutical Sciences, University of Iceland, Hagi, Hofsvallagata 53, IS-107 Reykjavik, Iceland; Department of Biological Sciences, University of Illinois at Chicago, 900 S Ashland Ave, Chicago IL 60607

**Keywords:** gut microbes, *in vitro* cultivation, insoluble fiber, natural products, zebrafish, metabolomics

## Abstract

There is a gap in measured microbial diversity when comparing genomic sequencing techniques versus cultivation from environmental samples in a laboratory setting. Standardized methods in artificial environments may not recapitulate the environmental conditions that native microbes require for optimal growth. For example, the intestinal tract houses microbes at various pH values as well as minimal oxygen and light environments. These microbes are also exposed to an atypical source of carbon: dietary fiber compacted in fecal matter. To investigate how the addition of insoluble fiber to isolation media could affect the cultivation of microbes from zebrafish intestines, an isolate library was built and analyzed using the bioinformatics pipeline IDBac. The addition of fiber led to an increase in bacterial growth and encouraged the growth of species from several phyla. Furthermore, fiber addition altered the metabolism of the cultivated gut-derived microbes and induced the production of unique metabolites that were not produced when microbes were otherwise grown on standard isolation media. Addition of this inexpensive carbon source to media supported the cultivation of a diverse community whose specialized metabolite production may more closely replicate their metabolite production *in vivo*.

## Introduction

Over the past 40 years, scientists searched for alternative strategies to increase the diversity of microbes isolated from environmental samples in a laboratory setting. Dubbed in 1932 as “the great plate count anomaly”, Razumov *et al.* noted that there was a large discrepancy between the viable plate count of aquatic bacteria on petri dishes compared to direct microscopic counts from the same environmental habitat.^1^ As more researchers reiterated the lack of a diverse community *in vitro*, confirming the phenomenon, a need for modifications in the cultivation process were imperative to improve isolation.

There are numerous variables that can be altered such as pH, salinity, temperature, oxygen levels, incubation time, and carbon and nitrogen substrates to create an *in vitro* environment that better represents the *in vivo* environment.^2^ Microbes have evolved to live in niche environments and their survival is dependent on the available simple or complex carbon and nitrogen substrates. Therefore, if a laboratory medium is lacking the main organic substrates that a microbe has evolved to use, this may help contribute to the plate count anomaly as the microbe may fail to grow *in vitro.* One of the earliest examples in altering the carbon substrate in a medium is the addition of the water-insoluble fraction of Alfalfa hay to petri dishes. This addition lead to an increase in colony counts of the microbes that colonize the rumen of cattle.^3^ Others contain substrates such as glass or cotton-fibers as an alternative for agar to evade fluctuating seaweed prices.^4,5^ Altering carbon sources gained further attention in the mid-2000’s with reports that modifying the organic carbon substrate in media recipes altered microbe growth and sporulation of environmental isolates.^6,7^ This catalyzed a variety of studies that altered media to increase bacterial production of pharmaceutically relevant molecular classes, escalate potential biodiesel sources, as well as encourage growth of specific bacterial species from animal microbiomes.^8–10^ This research solidified the foundation for what is now common practice to utilize several media when attempting to cultivate microbes from environmental samples (Figure S1). Recent studies have revealed that utilizing different organic carbon substrates can alter bacterial metabolism, leading to the discovery of novel compounds and potential biofilm inhibitors.^11,12^

Taking these developments into consideration, we were interested in cultivating the zebrafish gut microbiome to build a library of bacteria for testing in high-throughput antimicrobial and biofilm assays.^13,14^ When considering the native environment of vertebrate intestines, the main components of the system include the dietary nutrients entering the system for digestion and the indigestible nutrients that represent the fecal matter. Therefore, for microbes living in the intestinal tract, dietary fiber may serve as a large resource of nutrients and carbon. To optimize bacterial cultivation and isolation from zebrafish intestines, we utilized four types of media varying in nutrient density. Interestingly, the addition of organic insoluble fiber to the least nutrient medium led to an increase in the number of colonies observed on isolation plates as well as an increase in the diversity of colony morphologies.

Though the zebrafish gut microbiome varies slightly throughout development, once adulthood is reached, the gut microbiome is heavily conserved and strongly colonized by γ-Proteobacteria, Firmicutes, and Fusobacteria.^15,16^ Using the high-throughput bioinformatics pipeline IDBac,^17^ we isolated and analyzed 118 individual colonies from the zebrafish gut microbiome. IDBac analysis showed that our cultivation technique generated a taxonomically diverse library, reflecting that of previous metagenomic studies of the zebrafish gut microbiome, *vide supra*. Confirmation of a subset of the isolates with 16S rRNA gene sequencing showed that out of the 118 bacterial colonies isolated, our library represents bacterial isolates from three phyla. In this study we show that addition of insoluble fiber to low nutrient agar can increase the number of microbes isolated from vertebrate intestines. This simple, affordable addition also alters the specialized metabolite production of bacterial isolates allowing the bacteria to produce different metabolites that may be more similar to those produced *in vivo* in an *in vitro* environment.^18^

## Results

### Addition of insoluble fiber to low nutrient agar plates increases bacterial growth

To cultivate gut-derived microbes from zebrafish intestines, four types of agar media were used to target both fast and slow growing bacteria. Plates varied in nutrient density with high vitamin freshwater (HVF), NTF, SNF + fiber, and simple nutrient freshwater (SNF) decreasing in nutrient density respectively (**Figure 1**).^7,19^ Over 100 individual colonies were isolated over three months of growth to construct a zebrafish gut microbiome library. Addition of fiber to the least nutrient medium (SNF) led to a 1.7-fold increase in bacterial colony growth, surpassing the highest nutrient medium (HVF). Dissection of two healthy zebrafish intestines and two plating techniques in triplicate led to the isolation of 23, 38, 36, and 21 individual colonies from the HVF, NTF, SNF + fiber, and SNF plates, respectively.

**Figure 1:**
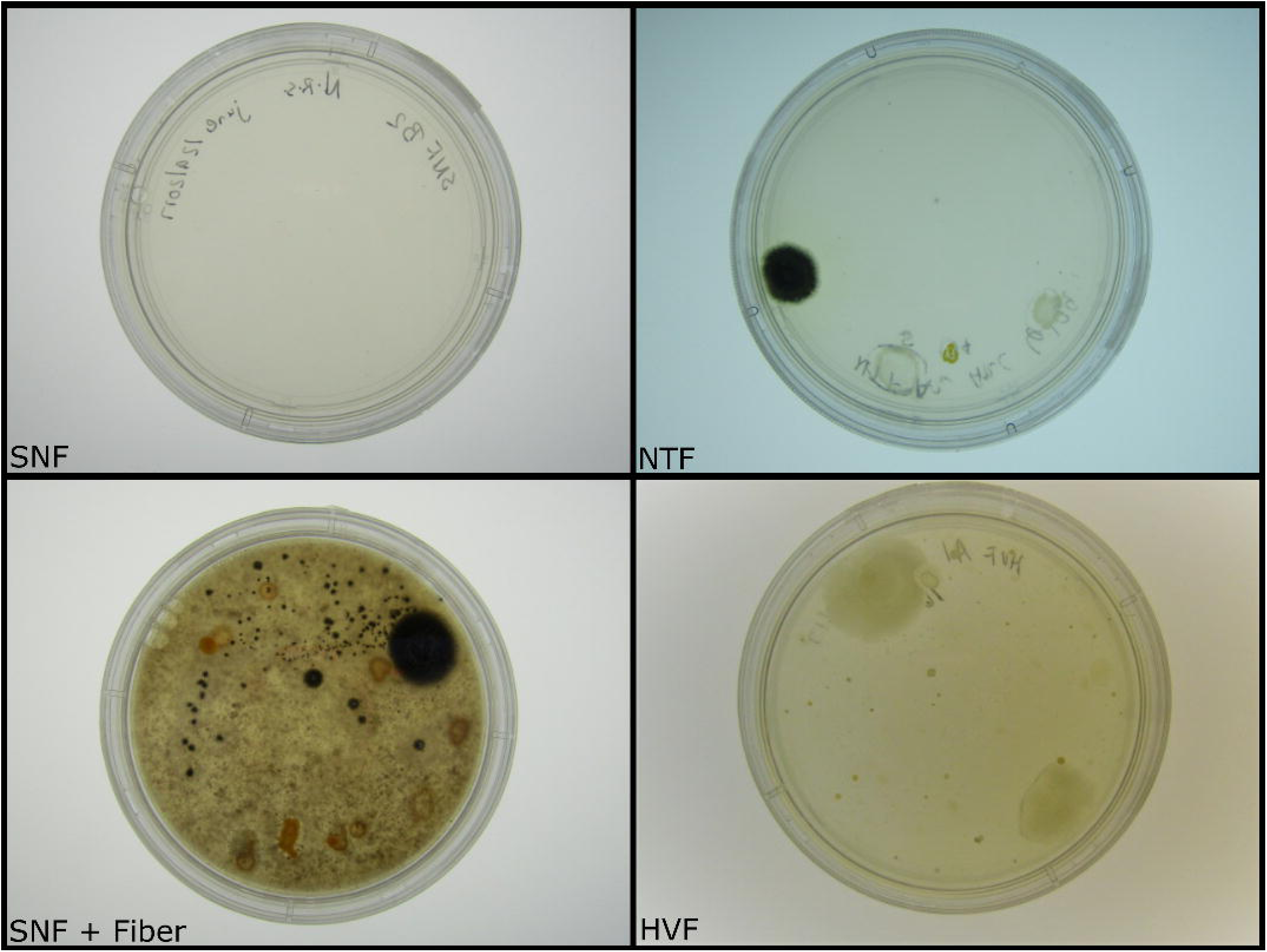
Diversity plates of isolated gut microbes. Using four types of agar, we observed a significant increase of bacterial growth in plates supplemented with insoluble fiber. The amount of growth observed in fiber supplemented plates (SNF + Fiber) was equivalent to that documented in high vitamin plates (HVF).

### IDBac analysis reveals that addition of insoluble fiber supports a diverse cultivation of gut microbes from agar plates

Library isolates were prepared for bioinformatics analysis using IDBac, a matrix-assisted laser desorption/ionization time-of-flight (MALDI-TOF) mass spectrometry pipeline that allows for rapid analysis of microbial proteins and specialized metabolites.^17,20^ The standard IDBac workflow uses a 70% aqueous formic acid solution to lyse cells but we found that this procedure was not efficient at lysing the more mucoid bacterial colonies isolated.^20^ A trifluoroacetic acid (TFA) extraction prior to plating samples for analysis was the most effective method to consistently collect viable MS fingerprint profiles for IDBac analysis.^21^ Measuring proteins in the 3,000 to 15,000 Da range, IDBac organized the microbial isolates into a pseudo-phylogeny dendrogram based on MS fingerprint protein similarity.^17^ **Figure 2** highlights 82 of the 118 bacterial isolates from our library. 16S rRNA sequencing was performed on a subset of isolates. In tandem, these analyses confirmed the generated library is diverse containing a minimum of 15 identified species within three phyla: Actinobacteria, Proteobacteria, and Firmicutes.

**Figure 2:**
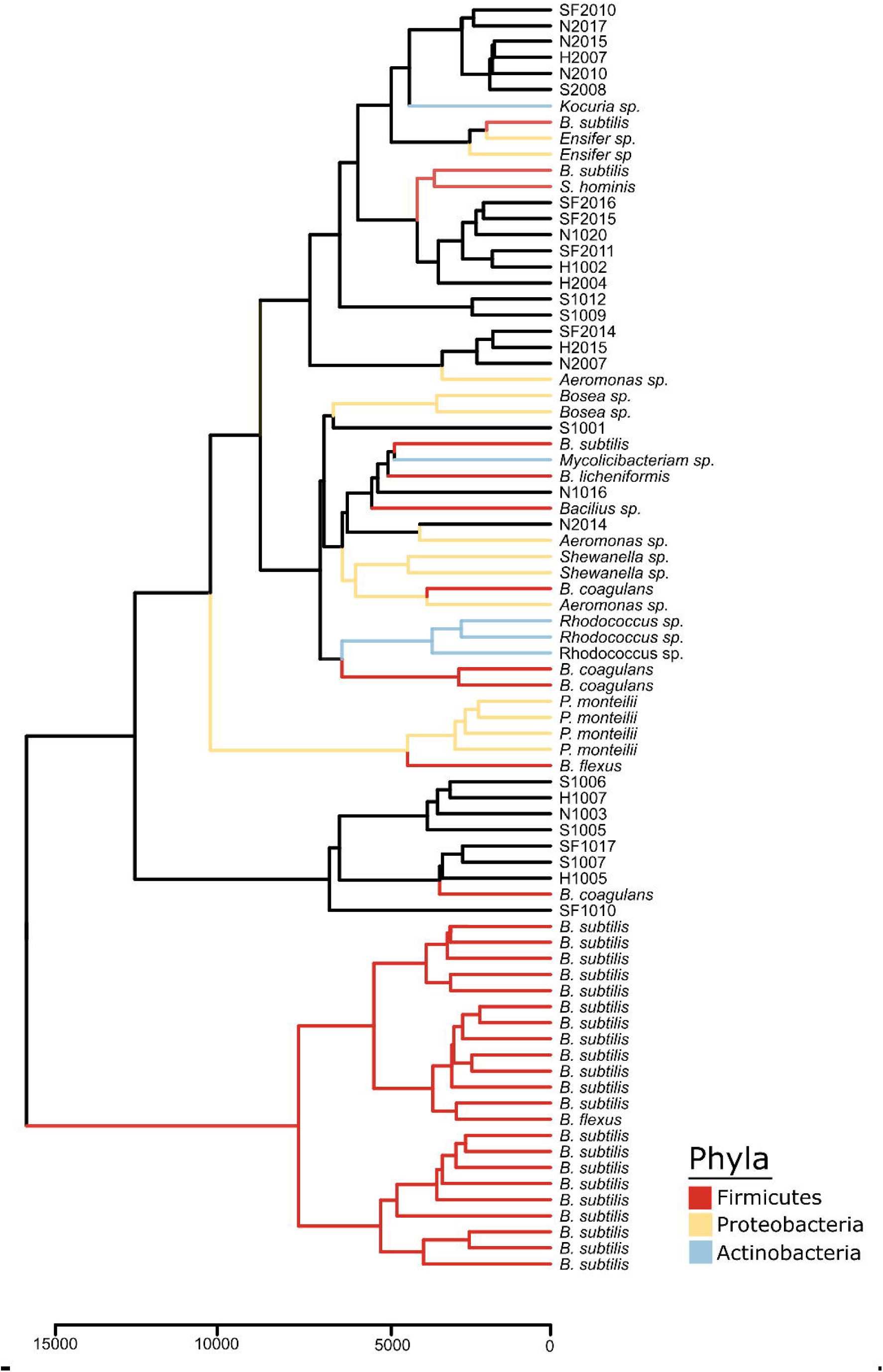
IDBac constructed dendrogram of cultivated bacterial isolates from the zebrafish gut. Our library is composed of over 80 isolates covering three phyla. To generate the IDBac dendrogram the following analysis settings were used: percent presence: 70, Signal to noise ratio: 4, Lower mass cutoff: 3,000, Upper mass cutoff: 15,0000, ppm tolerance: 1000. Distance algorithm: euclidean, Clustering algorithm: ward.D2, used presence/absence setting for dendrogram cladding.

### Metabolite association network highlights changes in metabolite production with different carbon sources

The IDBac pipeline also generates metabolite association networks (MANs) for rapid visualization of metabolite production and allows for a simple comparison between bacterial isolates grown in different conditions.^20^ **Figure 3A** shows a dendrogram highlighting 16 isolates that were grown on ISP2 agar, commonly used for lab cultivation, or SNF + fiber agar plates.^22^ In a MAN, nodes correspond to ions detected in a sample and the relationships between production across strains or conditions are visualized as arrows connecting nodes. The MANs of two isolates (**Figure 3B**; N2015 and S2008) imply that both have unique specialized metabolite production potential and when grown on agar supplemented with fiber, there is an increase in the unique specialized metabolites detected. The MAN for isolate SF2016 (**Figure 3C**), which was cultivated from a SNF + Fiber plate, is a robust producer of specialized metabolites. Interestingly, when comparing the two conditions, there is a larger number of nodes detected when grown without fiber.

**Figure 3:**
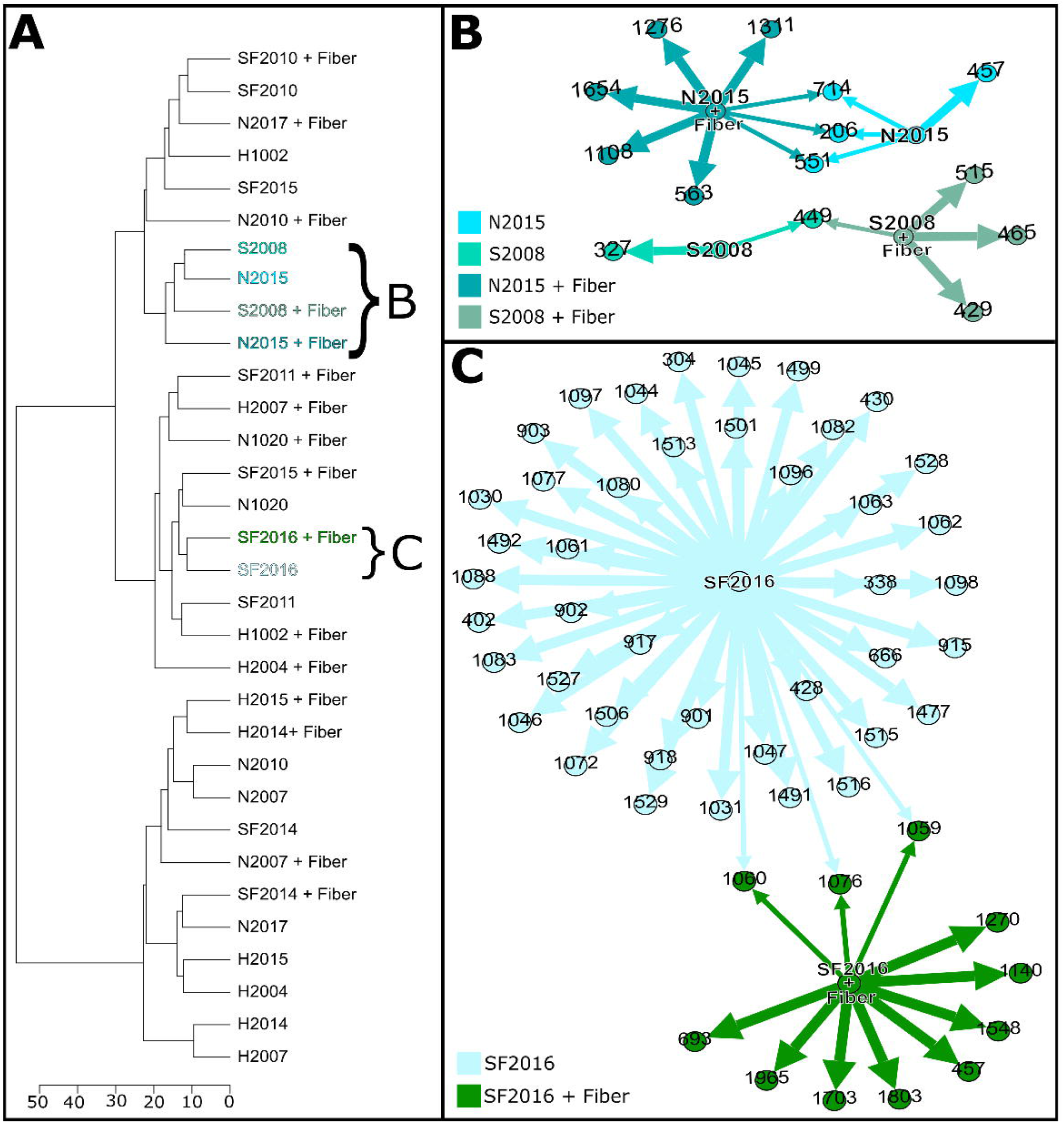
Comparison of metabolite production with and without fiber. (A) IDBac dendrogram of 16 isolates grown with or without fiber. (B) Metabolite association network (MAN) of isolates S2008 and N2015 grown with and without fiber. (C) MAN of isolate SF2016 comparing the metabolites produced when grown with or without fiber. Any *m/z* nodes corresponding to matrix or agar controls were subtracted prior to analysis. To generate the MANs the analysis settings used were as follows: percent presence: 70, Signal to noise:4, Lower mass cutoff: 200, Upper mass cutoff: 20000, ppm tolerance: 1000. Rather than displaying the MAN for the entire dendrogram, specific samples were highlighted and the corresponding MAN was downloaded to visualize in open graph platform Gephi (version 0.9.2) using the Yifan Hu layout.^44^

### Growth of gut microbes with insoluble fiber significantly increases the number of unique metabolites produced

High resolution liquid chromatography mass spectrometry (HR-LC-MS) of bacterial extracts from library isolates, SF2016, N2105, and S2008, were analyzed using XCMS, a metabolomic platform for statistical analyses of HR-LC-MS data. Through pairwise comparisons, XCMS can determine features (ion and retention time pair) that are up regulated in one extract versus another and visualize these differences via a cloud plot. As seen in **Figure 4A & B**, a cloud plot compares the total ion chromatograms (TIC) of two samples (i.e SF2016 grown on fiber agar vs. fiber agar control). Upregulated features are denoted by a circle and the darker the color of the circle, the more significant the signal. The cloud plot in **Figure 4A** highlights 49 features that are significantly upregulated by SF2016 when grown on fiber agar versus the 15 features observed when grown on non-fiber medium (p<0.001; **Figure 4B**). Similar cloud plots with the isolates N2015 and S2008 exhibit the increased trend with more features when the isolates were grown on fiber agar (p<0.001; **Figures S2 & S3**). Using XCMS meta-analysis, the number of significant features produced by the three isolates were organized into a Venn diagram to visualize the number of features unique to a specific isolate or shared between two or more (**Figure 4C & D**). When comparing the number of features produced by SF2016, N2015, and S2008 on fiber agar versus ISP2, the isolates had a 3.2, 3.3, and 1.5 fold change in features from growth with insoluble fiber.

**Figure 4:**
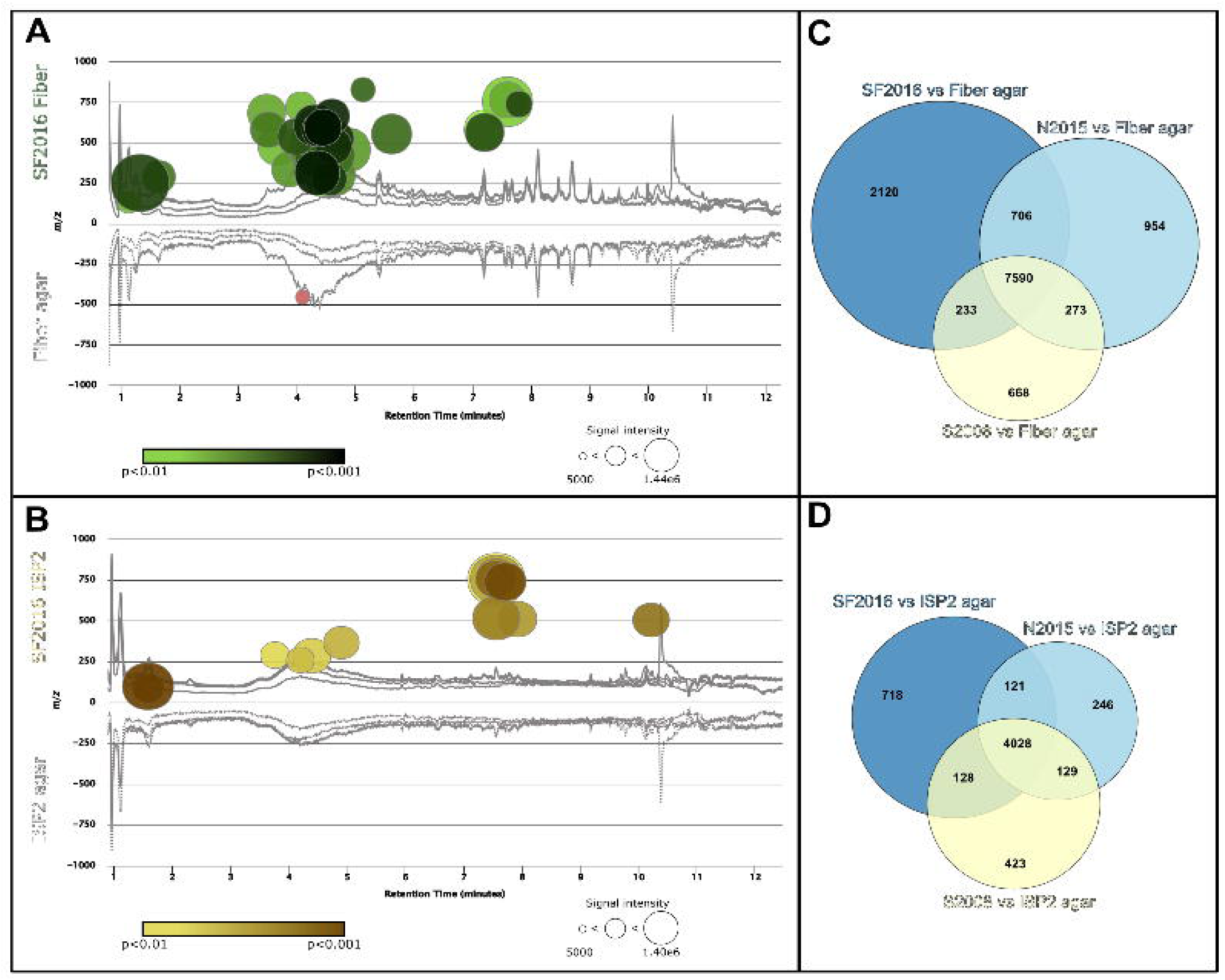
Gut microbe growth on fiber agar increases specialized metabolite production. HR-LC-MS data collected from metabolite extracts of library isolates SF2016, N2015, and S2008 were analyzed using XCMS pairwise and meta analysis. (A) Cloud plot comparison of SF2016 grown on fiber agar (top) and fiber agar control (bottom) highlighting significant metabolites (dots) produced by SF2016 when grown on fiber agar. The intensity of color of a circle correlates to the increased significance of the upregulated signal. (B) Cloud plot comparison of SF2016 grown on ISP2 agar (top) and ISP2 agar control (bottom). The Venn diagrams on the right represents the number of signals that are unique to each library isolate when grown on (C) fiber agar or (D) ISP2 medium.

## Discussion

Microbial libraries for high-throughput drug discovery screening are typically generated using two to six isolation media, varying in nutrient density, which select for fast or slow growing bacteria from an environmental sample.^23–26^ To generate a diverse library of gut-derived cultivable microbes from the zebrafish intestine as well as capture the unique metabolite production of this community, it is important to consider the carbon source(s) in the media. With dietary fiber being a highly abundant substrate for microbes colonizing the intestines, a fourth type of medium was utilized that incorporated the addition of insoluble fiber into SNF (the least nutrient medium). Using four types of media varying from high nutrient density to low, HVF, NTF, SNF + Fiber, and SNF respectively, 118 bacterial colonies were isolated to constitute a zebrafish gut microbiome library (**Figure 1**). The addition of wet-autoclaved insoluble fiber (20 g/L) lead to an increase in the growth of individual bacterial colonies (**Table S1**). The highest yield of bacterial isolates were collected from the average nutrient medium (NTF) however, the addition of fiber to the SNF lead to nearly an identical isolate yield (**Table S1**). Of note, the addition of fiber is a minimal and inexpensive addition compared to media typically used in environmental isolations (**Table S2**).

The potential taxonomic and specialized metabolite diversity of the cultivated isolates were assessed to confirm whether the isolates matched previous metagenomic sampling studies.^15,16^ To investigate the newly generated library, a bioinformatics platform known as IDBac was used to characterize the isolates. The IDBac workflow and software were designed to rapidly collect protein and specialized metabolite profiles of microbial libraries produced from environmental samples utilizing a MALDI-TOF mass spectrometer.^17,20^ Of the 118 isolates, 82 were examined using the IDBac workflow to organize the microbial library (**Figure 2**).The remaining 36 isolates were not analyzed due to either poor data quality collection during MALDI-TOF MS analysis or failure to recultivate from the initial glycerol stock (**Table S1**). Based on the MS protein fingerprints of the isolates, the newly constructed library represents three phyla. However, a number of isolates did not group into distinct branches nor with the IDBac in-house library that is built primarily from freshwater sponge-isolates,^17^ cheese microbes,^27^ and wild fish gut-derived microbes.^24^ Therefore, 16S rRNA sequencing on a subset of isolates in this library was performed to confirm the identity of a subset of the isolates and provide insight towards whether the cultivation efforts did support the growth of a diverse, gut microbe library (**Table S4**). Addition of taxonomic information gathered from 16S rRNA sequencing analysis to **Figure 2** supports the groupings formed from IDBac analysis.

Metagenomic studies of zebrafish intestines have shown that adults are colonized by a multitude of bacteria with γ-Proteobacteria, Firmicutes, and Fusobacteria as the most abundant members.^15,16^ 16S rRNA sequence analysis confirmed our cultivation strategy produced a library that accurately represents the major components of the zebrafish gut microbiome with representatives from the Firmicute, Actinobacteria, and Proteobacteria phyla (**Figure 2**). Bacteria from the three phyla have been linked to the gut and overall organism health. Firmicutes typically colonize the large intestines, primarily the colon, and are one of the most prevalent phyla represented in the gut microbiome.^28^ They have documented roles in fermentation of carbohydrates as well as lipid droplet formation for energy storage.^29,30^ In our dendrogram we observe the same trend with a considerable number of our isolates grouped into the Firmicute phylum (**Figure 2**). Of the Firmicutes isolated, *B. subtilis* was the most abundant isolate in this phylum. During its lifecycle in the gut, *B. subtilis* is known to form spores in half the time it takes for a laboratory isolate.^31^ Thus, it is possible the increase in isolation of *B. subtilis* in our library is due to the isolation of the more prolific, endogenous *B. subtilis* isolates from zebrafish intestines. *B. subtilis* has also been shown to protect fish from the aquatic pathogen *Aeromonas hydrophila*, which causes inflammation and steep mortality rates.^32^ With *A. hydrophila* infections being a persisting problem in zebrafish research facilities, the protective bioactivity of *B. subtilis* could contribute to the large representation of this species in our isolation efforts.^33^ Microbes responsible for maintaining homeostasis in the gut microbiome system fall under the Actinobacteria phylum.^34^ These maintenance microbes, which also reside in the colon, make up only a small portion of the gut microbiome community even though they hold a pivotal role in overall gut flora health.^28,34^ Once again, our dendrogram highlights the successful isolation and profiling of at least three genera of Actinobacteria such as *Rhodococcus, Kocuria*,and *Mycolicibacterium* (**Figure 2**).

Lastly, we observe Gram-negative Proteobacteria generally found in the small intestine. This population is minor with respect to the rest of the gut microbiome because many Proteobacteria are opportunistic and over colonization of these microbes has been strongly correlated with several human diseases.^35,36^ More specifically, we isolated and profiled microbes such as *Pseudomonas montelli, Aeromonas*, and *Shewanella* that have been previously isolated from feces and from aquatic environments respectively.^36,37^ No representatives from the Fusobacteria phylum were isolated because the cultivation methods used did not support the specific conditions needed to grow anaerobic bacteria. However, we hypothesize that due to the observed increase in cultivation of aerobic bacterial growth in this study, the same trend would hold if repeated with anaerobic conditions and would support the growth of Fusobacteria.

Following grouping by MS protein fingerprints with IDBac, specialized metabolite diversity was investigated using a metabolite association network (MAN) that corresponds to the IDBac generated dendrogram. Using IDBac, a dendrogram was created grouping 16 isolates grown on ISP2 or SNF + Fiber agar (**Figure 3A**). Selecting isolates N2015 and S2008 on the dendrogram generated two MANs that correspond to each of the isolates (**Figure 3B**). The MANs show signals found in both conditions(shared nodes) and unique signals (single nodes) that were detected when an isolate was grown with or without fiber. The MANs highlight that after removing nodes associated with matrix and the two types of mediums (controls), both isolates produce more unique specialized metabolites on SNF + Fiber than ISP2 (**Figure 3B**). Even though neither of these strains were originally isolated from SNF + Fiber plates but rather NTF and SNF media respectively, the MAN infers that metabolite production is increased by exposure to insoluble fiber during cultivation. Isolate SF2016 seems to be a prolific producer of unique chemistry based on the generated MAN (**Figure 3C**). Interestingly, the MAN suggests that SF2016 produces more unique specialized metabolites when grown on ISP2 medium rather fiber medium. In the case of SF2016, this is an example of how cultivation through the addition of insoluble fiber, allowed for the isolation of a prolific metabolite producer from the gut microbiome.

Having used IDBac as a prioritization tool, we aimed to further investigate and quantify the differences in metabolite production when gut microbes are grown on medium with insoluble fiber. Bacterial extracts of isolates SF2016, N2015, and S2008 grown on fiber or ISP2 agar, extracted, and analyzed via HR-LC-MS. The metabolomics platform XCMS was used to analyze and run statistical analysis on these extracts allowing for the selection of features that are significant (p<0.001) and abundant (based on fold change in intensities). XCMS can perform a suite of statistical analyses and output several visual representations of these statistical results with one example being cloud plots.^38^ **Figure 4A** shows a cloud plot comparing two TICs: SF2016 grown on fiber agar (top) and a fiber agar control (bottom). When grown on fiber agar SF2016 produces 41 features compared to 15 when SF2016 is grown on ISP2 (p<0.001; **Figure 4B**). If the significance threshold is relaxed to a p<0.01, the number of significant features between these two conditions are 174 and 49 respectively (**Figure S3**). Regardless of the threshold set for significance, SF2016 consistently makes 3-fold or more features when grown on medium supplemented with insoluble fiber than that without. The same trend holds with isolate N2015 (**Figure S4**) with an increase in unique metabolite production when grown on fiber agar. Interestingly, though S2008 does show unique metabolite production when grown on fiber agar, the cloud plot does not support that the addition of fiber lead to an increase in the number of metabolites produced, as observed in previous strains (**Figure S5**).

Using XCMS meta analysis, which was designed to prioritize interesting metabolite features from large metabolomic datasets, significant signals from each of the three isolates were identified and organized into a Venn diagram.^39^ Though SF2016 seems to be the most prolific of the three isolates when grown with fiber (**Figure 4C**), it is clear that the same trend holds in N2015 and S2008 with all three isolates showing an increase in metabolite production when grown on a medium with insoluble fiber compared to ISP2 (**Figure 4C & D**, respectively). Our HR-LC-MS results support that the addition of insoluble fiber to medium leads to an increase in specialized metabolite production in a natural product laboratory workflow.

It is worth noting that there are differences in the number of observed specialized metabolites between the IDBac MANs and the XCMS chromatograms. The differences in ions observed can arise for a variety of reasons. Importantly, these are two different ionization sources (MALDI vs electrospray ionization - ESI) and the exact mechanism by which ions are produced is fundamentally different. Various functional groups on the specialized metabolites themselves may favor one ionization method over the other and neither method is capable of comprehensively ionizing every metabolite in a given extract. Secondly, the samples subjected to the analyses are also fundamentally different. MALDI is incredibly fast (milliseconds per sample) and utilizes a small amount of a microbial colony, this direct detection typically favors singularly charged ions with little in-source fragmentation. Whereas ESI-LC-MS allows for a concentration of the colony and surrounding agar, which likely captures a better sampling of specialized metabolite production. Additionally, LC-MS analyses are time consuming with each run taking at least 12 mins/colony extraction. This allows for separation of different isomers which are not possible to distinguish in MALDI-TOF MS. However, ESI is known to give rise to in-source fragments which may falsely increase the perceived number of specialized metabolites in a given sample. All in all, while the overall numbers of specialized metabolites are different, the trends hold across these orthogonal methods. So the most prolific producer observed via the IDBac generated MAN yields the highest number of features in the ESI-LC-MS analysis.

One of the biggest challenges the field of natural products faces is the frequent rediscovery of known small molecules from environmental samples.^40^ Since the discovery of penicillin over 75 years ago, workflows which include cultivation of select genera mainly based on morphological features or 16S rRNA followed by bioassay guided fractionation have been used to continue searching for new drug candidates and/or scaffolds.^41^ This workflow has been historically successful in elucidating novel specialized metabolites produced in high titer by environmental microbes. However, advances in technologies such as bioinformatics, mass spectrometry, proteomics, and synthetic biology have improved our ability to identify bioactive metabolites that are produced in smaller titers. A distinct limitation of these methods however, is their inability to detect or predict novel metabolites while assessing taxonomic diversity.^41^ Therefore, it is important to consider how changes in cultivation techniques, such as altering carbon sources, can alter the isolation of specific microbes as well as the abundance and variety of metabolites produced. As discussed in the introduction, work has already been completed showing that altering nutrients or growth conditions can lead to increases in cultivation, altered specialized metabolism, and the discovery of putative drug candidates. Thus, developing new methods to propagate microbial metabolite production prior to beginning natural product discovery efforts could be vital to assisting future drug discovery.

With the goal of constructing a taxonomically and chemically diverse zebrafish gut microbe library, insoluble fiber was added to isolation medium as a carbon source. By providing this solid substrate during *in vitro* growth, we observed an increase in bacterial growth to isolate microbes from environmental samples. Harnessing the speed, accuracy, and sensitivity of the IDBac workflow, protein and metabolite profiles of the generated library supported that the increase in cultivation observed from the addition of fiber encouraged growth of the gut microbial community and suggested that providing insoluble fiber as a carbon source alters metabolite production of these isolates. Metabolomic statistical analysis of three bacterial isolates, SF2016, N2015, and S2008, confirmed that regardless of whether an isolate was originally cultivated from a SNF + Fiber plate, addition of insoluble fiber to growth medium lead to an increase in metabolite production. Further analysis is required to determine if the increase in metabolite production observed represents novel metabolites that have not been cultivated *in vitro* or if this cultivation technique allows for metabolite isolation that has traditionally only been observed *in vivo.* Work is ongoing to perform biological assays and identify constituents from the cultivable zebrafish gut microbes with antimicrobial and biofilm inhibition activity.

## Materials and Methods

### Agar plate preparation

Agar plates were prepared by following the medium recipes detailed in **Table S4**. Mediums were autoclaved for 45 minutes and cooled to 55°C prior to the addition of 25 mg/L of 0.22 micron sterile filtered nalidixic acid and cycloheximide (Sigma).

### Dissection of Zebrafish Intestines

Animals were treated and cared for in accordance with the National Institutes of Health Guide for the Care and Use of Laboratory animals. All experiments were approved by the University of Illinois at Chicago Institutional Animal Care and Use Committee (IACUC). Two adult *casper* zebrafish (lacking pigmentation),^42^ were sacrificed at >90 days post fertilization with 0.1% tricaine solution for 10 minutes and bathed in 70% EtOH prior to dissection. To dissect out intestines, a ventral incision was made from the front fin to the back fin. If the fish was female, eggs were separated and removed to expose the intestines. Both ends of the intestinal tract were cut and the tract was removed.

### Intestinal sample plating

The intestines were transferred to a 1x PBS solution with three sterilized glass beads. This solution was vortexed for several minutes to break up the intestinal tissues. A 10-fold dilution of the intestinal solution was made before proceeding. Intestinal solution was plated in triplicate using two plating techniques (1) spreading or (2) serial spotting. For (1) spreading, 100 *µ*L of the intestinal sample was pipetted to the agar and spread for even distribution of the sample. For serial spotting (2), a sterilized cotton swab was dipped into the intestinal solution and spotted throughout the plate. The entire plating procedure was then repeated with the 10-fold dilution. Plates were left to grow for up to three months at room temperature (∼25°C).

### Gut microbe isolation

Diversity plates were frequently monitored for new growth. New colonies were transferred to ISP2 to purify to single colonies. Once the purity of the culture was confirmed, a single colony was transferred to a liquid culture of ISP2 media. This liquid culture was left to shake at 225 RPM at 25°C until the liquid culture was turbid. Once turbid, the liquid culture was used to plate the isolate on ISP2 agar and frozen stocked.

### IDBac sample preparation

From the pure isolates grown on ISP2 or SNF + fiber media, a sterilized toothpick was used to transfer a portion of a bacterial colony to a sterilized microcentrifuge tube containing 5 *µ*L of HPLC grade TFA. Samples remained in TFA for 30 minutes to facilitate cell lysis and protein extraction. Following this incubation time, DI water (20 *µ*L) and acetonitrile (30 *µ*L) were added to each sample, briefly vortexed, and centrifuged at 10,000 rpm for two minutes. The supernatant (1.5 *µ*L) was mixed with 1.5 *µ*L of □-cyano-4-hydroxycinnamic acid (CHCA) matrix for a 1:1 ratio of sample to matrix. The 1:1 sample:matrix mixture was then plated on a MALDI target plate in technical triplicate for IDBac analysis (1 *µ*L). Measurements were performed in positive linear and reflection modes on an Autoflex Speed LRF mass spectrometer (Bruker Daltonics) equipped with a smartbeam-II laser (355 nm).

### 16S sequencing molecular analysis

Total genomic DNA was extracted using a commercially available DNeasy Ultraclean Microbial Kit (Qiagen) following the manufacturer’s recommendations. Primer pair 27F (5’-CAGAGTTTGATCCTGGCT-3’) and 1492R (5’-AGGAGGTGATCCAGCCGCA-3’) were used to amplify the partial 16S ribosomal RNA (rRNA) gene sequence.^43^ PCR conditions in a BioRad thermocycler were as follows: initial denaturation at 95°C for 5 minutes, followed by 35 cycles of denaturation at 95°C for 15 seconds, annealing at 60°C for 15 seconds, and extension at 72 °C for 30 seconds, and a final extension step at 72 °C for 2 minutes. Each amplified product was purified using QIAquick PCR Purification kit from Qiagen, followed by Sanger sequencing. Data analyzed by Geneious V11.1.4 software.

### IDBac protein analysis

To generate dendrograms, MALDI-TOF MS protein data was uploaded to IDBac. The following settings were used to generate reported dendrogram: percent presence: 70, signal to noise ratio: 4, lower mass cutoff: 3,000, upper mass cutoff: 15,000, ppm tolerance: 1000, distance algorithm: euclidean, clustering algorithm: ward.D2, used peak presence/absence as grouping criteria.

### IDBac Metabolite association networks

A new dendrogram was constructed in IDBac using MALDI-TOF MS data collected on the 16 isolates that were grown on both fiber and ISP2 medium using the same settings as previously. For MANs, isolates SF2016, N2015 and S2008 were selected and the following settings were applied; percent presence: 70, signal to noise ratio: 4, lower mass cutoff: 200, upper mass cutoff: 2,000, ppm tolerance: 1,000, sample subtraction: Matrix. The network data was downloaded as a .csv file from IDBac and importanted into Gephi version 0.9.2. This was repeated with sample subtraction being changed to fiber and ISP2 agar controls. In Gephi, MANs for the same isolate were compared and nodes corresponding to matrix or agar controls were manually removed to generate reported figures.

### Metabolite extraction from bacterial isolates

Isolates SF2016, N2105, and S2008 were plated on thin (3 mL of agar in 60 mm petri dish) SNF + Fiber or ISP2 media in duplicate. Samples and control plates were grown for three days at 25 °C. Entire contents of petri dish extracted with 3 mL of MeOH and sonicated for 30 minutes. Crude extract were used for HR-LC-MS analysis and stored at -20 °C. All extractions were performed in biological replicates (N =3).

### HR-LC-MS data collection and XCMS analysis

Biological extracts were pooled (1mg/mL) and run on a Bruker Impact II qTOF with a C18 UPLC (Phenomenex) in two technical replicates. The method for MS^1^ data collection was as follows: solvent A (H20 with 0.1% formic acid) and solvent B (ACN with 0.1% formic acid). 10% isocratic solvent B for 2 minutes, 10%-100% gradient of solvent B for 8 minutes, 2 minute wash of 100% solvent B and 2 minute equilibration. Data was converted to .mzXML from Bruker software Data Analysis and uploaded to the online metabolomics platform XCMS. Pairwise comparisons of SF2016F vs Fiber agar, SF2016I vs ISP2 agar, and SF2016F vs SF2016I were run using Parameter ID: UPLC/Bruker Q-TOF pos. The same analysis was run for isolates N2015 and S2008. Cloud plots were generated with the following settings; p-value: 0-0.001, fold change: ≥1.5, retention time: 1-12, intensity: ≥5,000. For XCMS meta analysis, pairwise jobs of each of the three isolates versus the fiber control were selected and run with the following settings; fold change ≥ 1.5, max p-value ≤ 0.001, max intensity ≥5,000, *m/z* tolerance 0.01, RT tolerance 60 secs. The same workflow was repeated for meta analysis of each isolate with ISP2 control.

## Acknowledgements

This publication was supported by the National Institute of General Medical Sciences of the National Institutes of Health under Award Number R01GM125943 (LMS and BTM); University of Illinois at Chicago Startup Funds (LMS); Icelandic Research Fund Grant 152336-051 (B.T.M. and Dr. Sesselja Omarsdottir); American Society for Pharmacognosy research startup grant (LMS). ARC was supported in part by the National Science Foundation Illinois Louis Stokes Alliance for Minority Participation Bridge to the Doctorate Fellowship (grant number 1500368) and a UIC Abraham Lincoln retention fellowship.

## References: No more than 85

1. Staley JT, Konopka A. Measurement of in situ activities of nonphotosynthetic microorganisms in aquatic and terrestrial habitats. Annu Rev Microbiol 1985; 39:321–46.

2. Lagier J-C, Edouard S, Pagnier I, Mediannikov O, Drancourt M, Raoult D. Current and past strategies for bacterial culture in clinical microbiology. Clin Microbiol Rev 2015; 28:208–36.

3. Chung KT, Hungate RE. Effect of alfalfa fiber substrate on culture counts of rumen bacteria. Appl Environ Microbiol 1976; 32:649–52.

4. Moraes-Cerdeira RM, Krans JV, McChesney JD, Pereira AMS, Franca SC. Cotton Fiber as a Substitute for Agar Support in Tissue Culture. HortScience 1995; 30:1082–3.

5. Ferris MJ, Hirsch CF. Method for isolation and purification of cyanobacteria. Appl Environ Microbiol 1991; 57:1448–52.

6. Zhao S, Shamoun SF. The effects of culture media, solid substrates, and relative humidity on growth, sporulation and conidial discharge of Valdensinia heterodoxa. Mycol Res 2006; 110:1340–6.

7. Jensen PR, Gontang E, Mafnas C, Mincer TJ, Fenical W. Culturable marine actinomycete diversity from tropical Pacific Ocean sediments. Environ Microbiol 2005; 7:1039–48.

8. Tran H, Stephenson S, Pollock E. Evaluation of Physarum polycephalum plasmodial growth and lipid production using rice bran as a carbon source. BMC Biotechnol 2015; 15:67.

9. Varghese LM, Agrawal S, Sharma D, Mandhan RP, Mahajan R. Cost-effective screening and isolation of xylano-cellulolytic positive microbes from termite gut and termitarium. 3 Biotech 2017; 7:108.

10. Bisht D, Yadav SK, Gautam P, Darmwal NS. Simultaneous production of alkaline lipase and protease by antibiotic and heavy metal tolerant Pseudomonas aeruginosa. J Basic Microbiol 2013; 53:715–22.

11. Wang B, You J, King JB, Cai S, Park E, Powell DR, Cichewicz RH. Polyketide glycosides from Bionectria ochroleuca inhibit Candida albicans biofilm formation. J Nat Prod 2014; 77:2273–9.

12. Timmermans ML, Picott KJ, Ucciferri L, Ross AC. Culturing marine bacteria from the genus Pseudoalteromonas on a cotton scaffold alters secondary metabolite production. Microbiologyopen 2018; :e00724.

13. Peach KC, Bray WM, Shikuma NJ, Gassner NC, Lokey RS, Yildiz FH, Linington RG. An image-based 384-well high-throughput screening method for the discovery of biofilm inhibitors in Vibrio cholerae. Mol Biosyst 2011; 7:1176–84.

14. Wong WR, Oliver AG, Linington RG. Development of antibiotic activity profile screening for the classification and discovery of natural product antibiotics. Chem Biol 2012; 19:1483–95.

15. Stephens WZ, Burns AR, Stagaman K, Wong S, Rawls JF, Guillemin K, Bohannan BJM. The composition of the zebrafish intestinal microbial community varies across development. ISME J 2016; 10:644–54.

16. Roeselers G, Mittge EK, Stephens WZ, Parichy DM, Cavanaugh CM, Guillemin K, Rawls JF. Evidence for a core gut microbiota in the zebrafish. ISME J 2011; 5:1595–608.

17. Clark CM, Costa MS, Sanchez LM, Murphy BT. Coupling MALDI-TOF mass spectrometry protein and specialized metabolite analyses to rapidly discriminate bacterial function. Proc Natl Acad Sci U S A 2018; 115:4981–6.

18. Davies J. Specialized microbial metabolites: functions and origins. J Antibiot 2013; 66:361–4.

19. Hong K, Gao A-H, Xie Q-Y, Gao H, Zhuang L, Lin H-P, Yu H-P, Li J, Yao X-S, Goodfellow M, et al. Actinomycetes for marine drug discovery isolated from mangrove soils and plants in China. Mar Drugs 2009; 7:24–44.

20. Clark CM, Costa MS, Conley E, Li E, Sanchez LM, Murphy BT. Using the Open-Source MALDI TOF-MS IDBac Pipeline for Analysis of Microbial Protein and Specialized Metabolite Data. J Vis Exp [Internet] 2019; Available from: http://dx.doi.org/10.3791/59219

21. Freiwald A, Sauer S. Phylogenetic classification and identification of bacteria by mass spectrometry. Nat Protoc 2009; 4:732–42.

22. Mohseni M, Norouzi H, Hamedi J, Roohi A. Screening of antibacterial producing actinomycetes from sediments of the caspian sea. Int J Mol Cell Med 2013; 2:64–71.

23. Ochoa JL, Sanchez LM, Koo B-M, Doherty JS, Rajendram M, Huang KC, Gross CA, Linington RG. Marine Mammal Microbiota Yields Novel Antibiotic with Potent Activity Against Clostridium difficile. ACS Infect Dis 2018; 4:59–67.

24. Sanchez LM, Wong WR, Riener RM, Schulze CJ, Linington RG. Examining the fish microbiome: vertebrate-derived bacteria as an environmental niche for the discovery of unique marine natural products. PLoS One 2012; 7:e35398.

25. Mincer TJ, Jensen PR, Kauffman CA, Fenical W. Widespread and persistent populations of a major new marine actinomycete taxon in ocean sediments. Appl Environ Microbiol 2002; 68:5005–11.

26. Elfeki M, Alanjary M, Green SJ, Ziemert N, Murphy BT. Assessing the Efficiency of Cultivation Techniques To Recover Natural Product Biosynthetic Gene Populations from Sediment. ACS Chem Biol 2018; 13:2074–81.

27. Wolfe BE, Button JE, Santarelli M, Dutton RJ. Cheese rind communities provide tractable systems for in situ and in vitro studies of microbial diversity. Cell 2014; 158:422–33.

28. Jandhyala SM, Talukdar R, Subramanyam C, Vuyyuru H, Sasikala M, Nageshwar Reddy D. Role of the normal gut microbiota. World J Gastroenterol 2015; 21:8787–803.

29. Huang Y, Shi X, Li Z, Shen Y, Shi X, Wang L, Li G, Yuan Y, Wang J, Zhang Y, et al. Possible association of Firmicutes in the gut microbiota of patients with major depressive disorder. Neuropsychiatr Dis Treat 2018; 14:3329–37.

30. Duncan SH, Louis P, Flint HJ. Cultivable bacterial diversity from the human colon. Lett Appl Microbiol 2007; 44:343–50.

31. Tam NKM, Uyen NQ, Hong HA, Duc LH, Hoa TT, Serra CR, Henriques AO, Cutting SM. The intestinal life cycle of Bacillus subtilis and close relatives. J Bacteriol 2006; 188:2692–700.

32. Kong W, Huang C, Tang Y, Zhang D, Wu Z, Chen X. Effect of Bacillus subtilis on Aeromonas hydrophila-induced intestinal mucosal barrier function damage and inflammation in grass carp (Ctenopharyngodon idella). Sci Rep 2017; 7:1588.

33. Mocho J-P, Martin DJ, Millington ME, Saavedra Torres Y. Environmental Screening of Aeromonas hydrophila, Mycobacterium spp., and Pseudocapillaria tomentosa in Zebrafish Systems. J Vis Exp [Internet] 2017; Available from: http://dx.doi.org/10.3791/55306

34. Binda C, Lopetuso LR, Rizzatti G, Gibiino G, Cennamo V, Gasbarrini A. Actinobacteria: A relevant minority for the maintenance of gut homeostasis. Dig Liver Dis 2018; 50:421–8.

35. Sharma KK, Kalawat U. Emerging infections: shewanella - a series of five cases. J Lab Physicians 2010; 2:61–5.

36. de Abreu PM, Farias PG, Paiva GS, Almeida AM, Morais PV. Persistence of microbial communities including Pseudomonas aeruginosa in a hospital environment: a potential health hazard. BMC Microbiol 2014; 14:118.

37. Dailey FE, McGraw JE, Jensen BJ, Bishop SS, Lokken JP, Dorff KJ, Ripley MP, Munro JB. The Microbiota of Freshwater Fish and Freshwater Niches Contain Omega-3 Fatty Acid-Producing Shewanella Species. Appl Environ Microbiol 2016; 82:218–31.

38. Tautenhahn R, Patti GJ, Rinehart D, Siuzdak G. XCMS Online: a web-based platform to process untargeted metabolomic data. Anal Chem 2012; 84:5035–9.

39. Patti GJ, Tautenhahn R, Siuzdak G. Meta-analysis of untargeted metabolomic data from multiple profiling experiments. Nat Protoc 2012; 7:508–16.

40. Pye CR, Bertin MJ, Lokey RS, Gerwick WH, Linington RG. Retrospective analysis of natural products provides insights for future discovery trends. Proc Natl Acad Sci U S A 2017; 114:5601–6.

41. Katz L, Baltz RH. Natural product discovery: past, present, and future. J Ind Microbiol Biotechnol 2016; 43:155–76.

42. White RM, Sessa A, Burke C, Bowman T, LeBlanc J, Ceol C, Bourque C, Dovey M, Goessling W, Burns CE, et al. Transparent adult zebrafish as a tool for in vivo transplantation analysis. Cell Stem Cell 2008; 2:183–9.

43. Lane DJ. 16S/23S rRNA sequencing.(eds. Stackebrandt E. Goodfellow M) Nucleic Acid Tech. Bact. Syst. 115--175. 1991;

44. Hu Y. Efficient, high-quality force-directed graph drawing. Mathematica Journal 2005; 10:37–71.

