## Supplemental Info for "Addition of insoluble fiber to isolation media allows for increased metabolite diversity of lab-cultivable microbes derived from zebrafish gut samples"

| Figure S1: Microbial carbon sources………………………………………………………… | S2 |
| --- | --- |
| Figure S2: MAN of isolate SF2010…………………………………………………………… | S3 |
| Figure S3: XCMS cloud plot comparison of SF2016 extracts (p<0.01)………………….. | S4 |
| Figure S4: XCMS cloud plot comparison of N2015 extracts (p<0.001)........................... | S5 |
| Figure S5: XCMS cloud plot comparison of S2008 extract (p<0.001)………………….... | S6 |
| Table S1: Library isolates and quality of data collection……………………….………….. | S7 |
| Table S2: Medium recipes…………………………………………………………………….. | S8 |
| Table S3: Results from 16S rRNA sequencing and accession numbers......................... | S9 |
| Data Availability……………………………………………………………………………….... | S13 |

Figure S1: Microbial carbon sources. Chemical structures of fiber components used in previous studies to increase microbial growth and/or metabolite production.

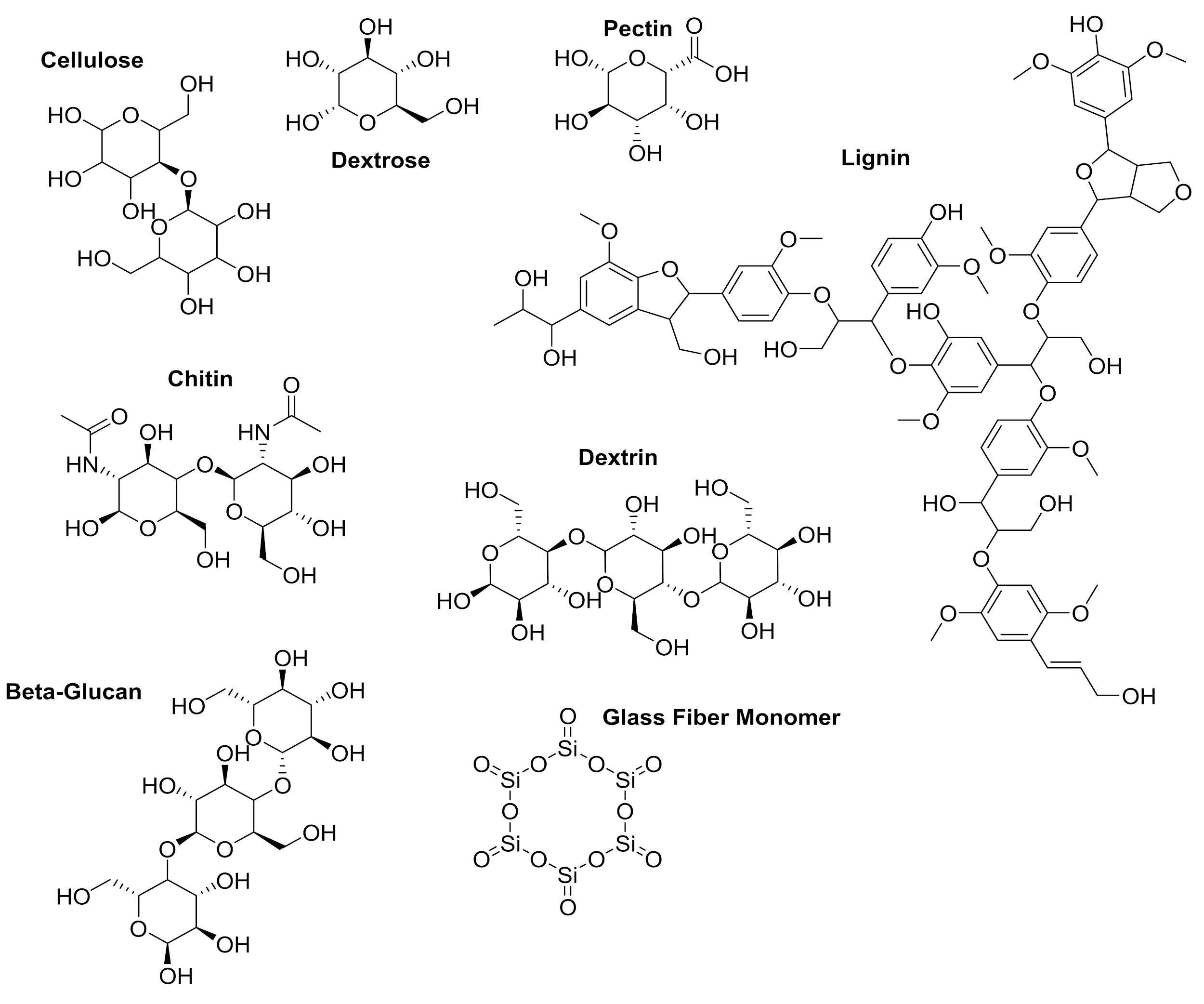

Figure S2: MAN of isolate SF2010. MAN suggests that isolate SF2010 produces unique chemistry when grown with insoluble fiber.

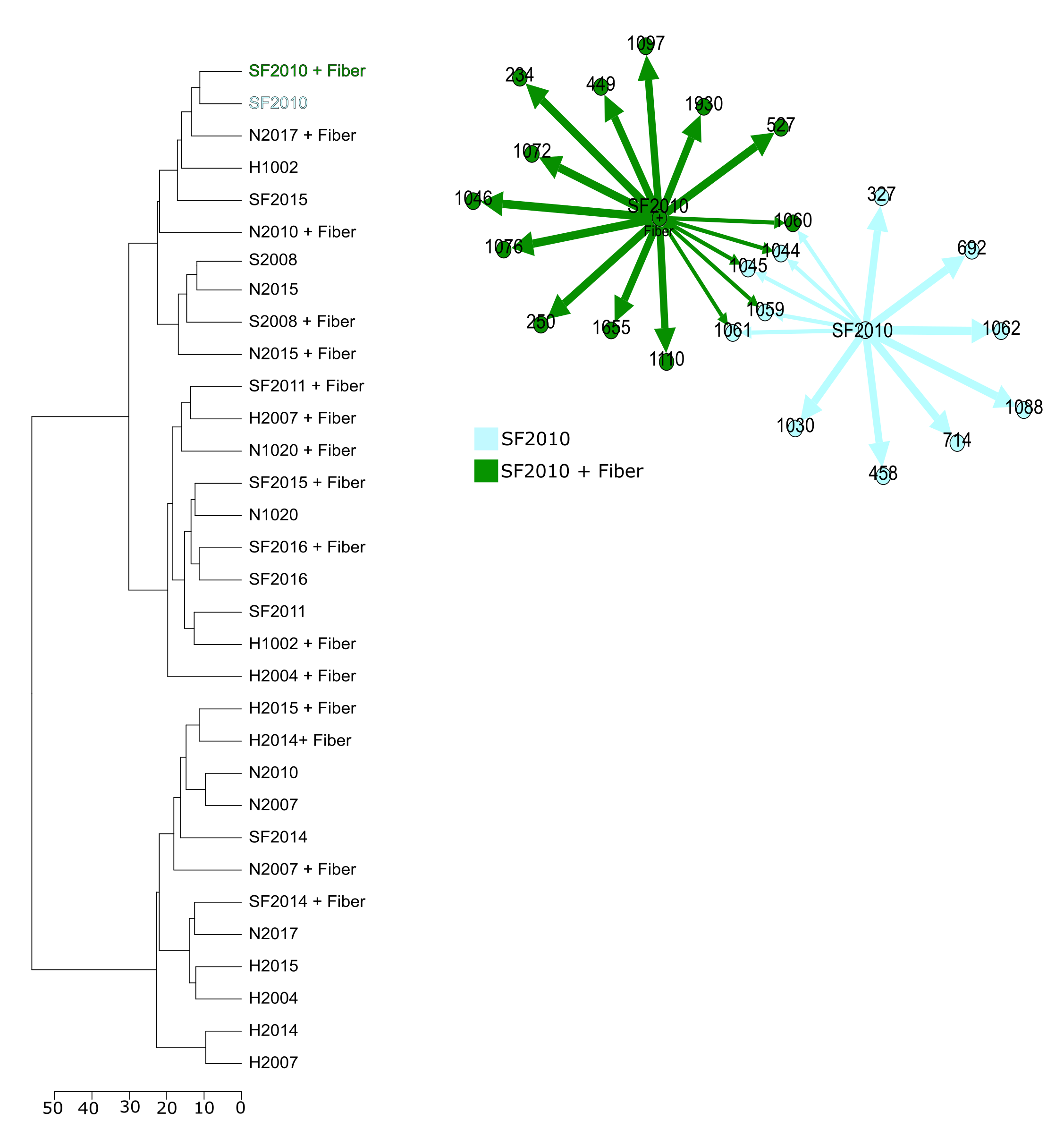

Figure S3: XCMS cloud plot comparison of SF2016 extracts. Metabolite production from SF2016 grown on fiber and compared with fiber agar control. (A) 174 metabolites were identified to be significantly upregulated by SF2016 when grown on fiber compared to (B) 49 metabolites on ISP2 (p<0.01).

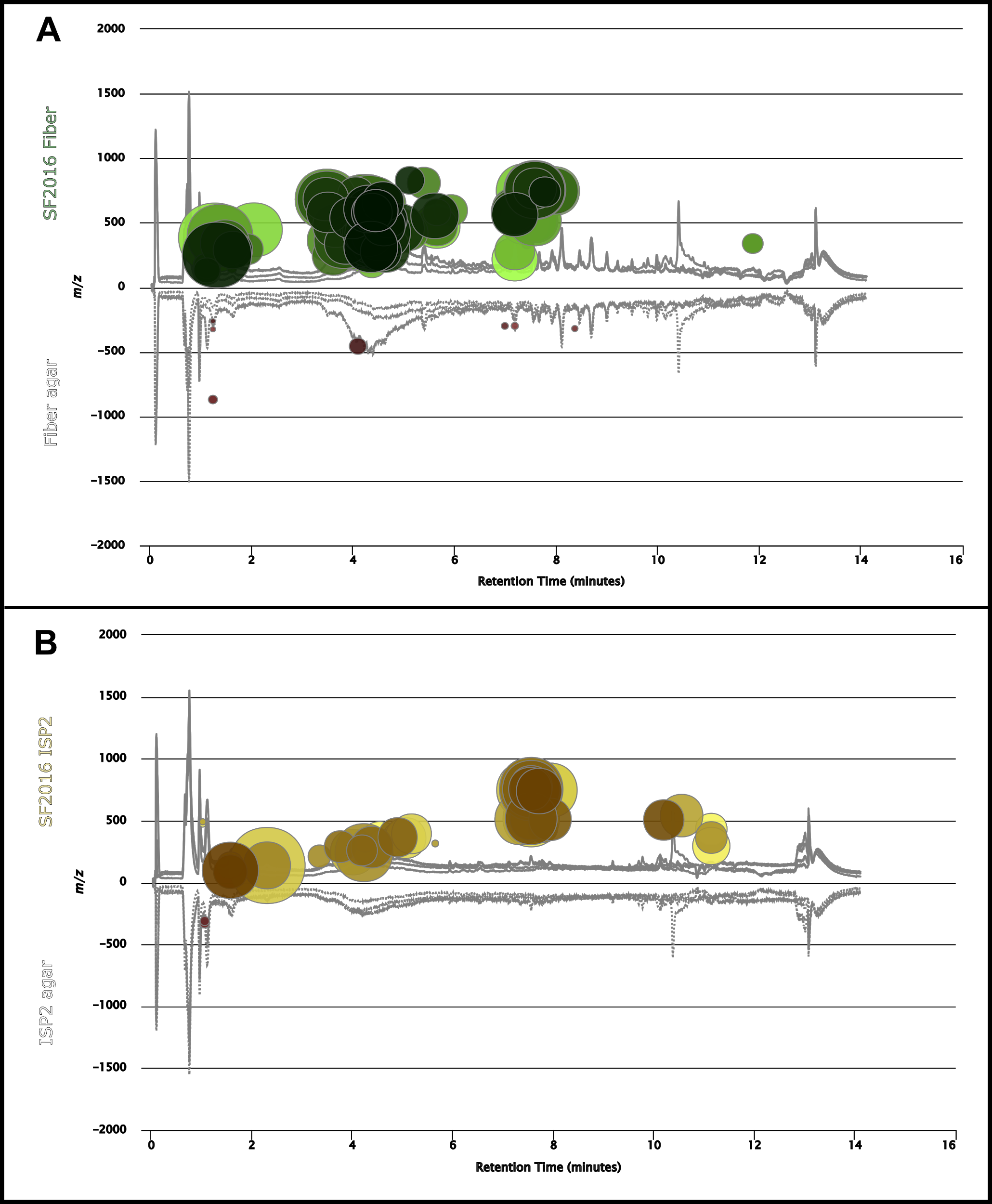

Figure S4: XCMS cloud plot comparison of N2015 extract. Metabolite production from N2105 grown on fiber and compared with fiber agar control. (A) 32 metabolites were identified to be significantly upregulated by N2015 (green) and 6 downregulated (red) when grown on fiber compared to (B) 1 upregulated metabolite on ISP2 (p<0.001).

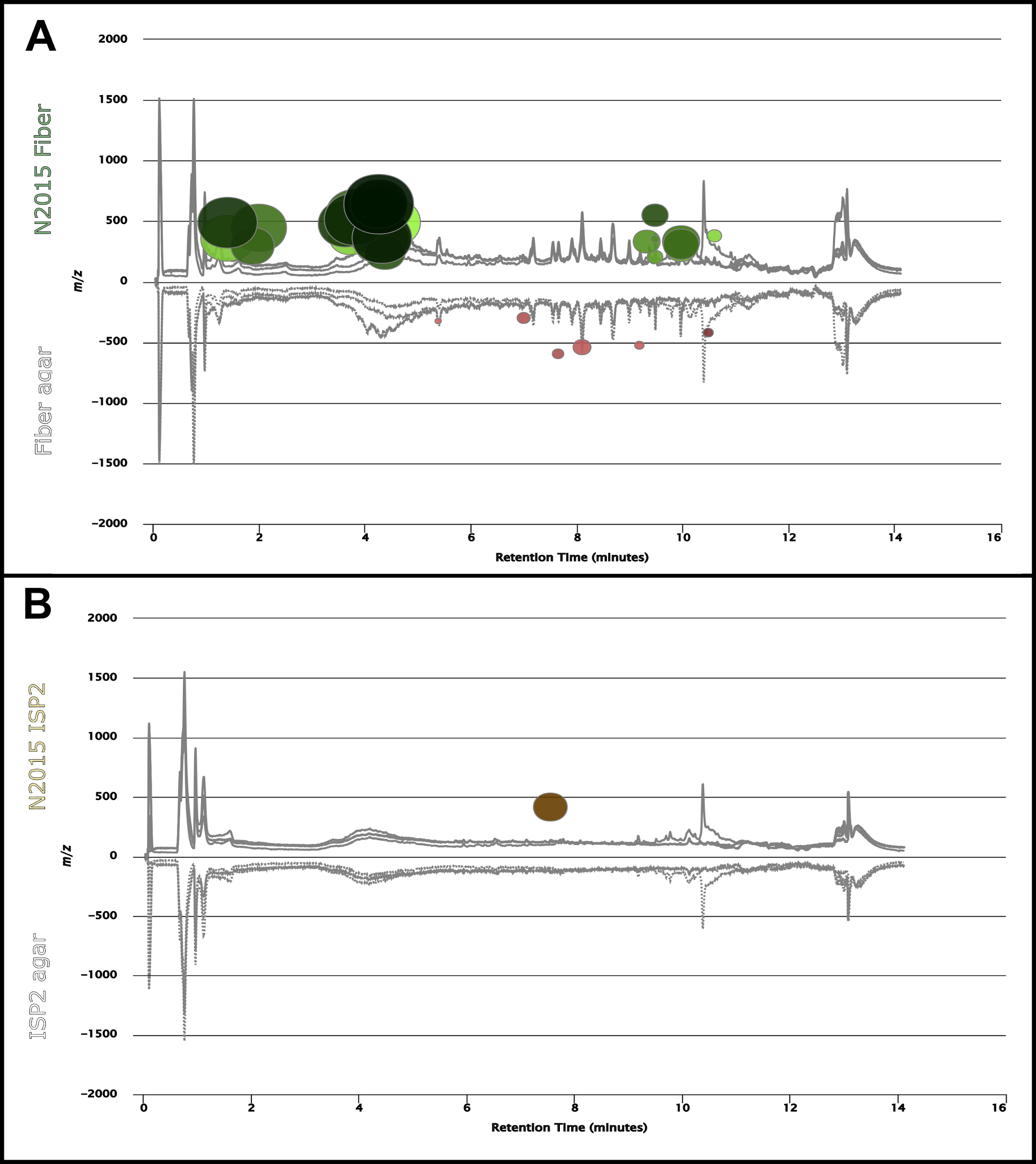

Figure S5: XCMS cloud plot comparison of S2008 extract. Metabolite production from S2008 grown on fiber and compared with fiber agar control. (A) 6 metabolites were identified to be significantly upregulated (green) and 2 downregulated (red) by S2008 when grown on fiber compared to (B) 9 metabolites that are upregulated on ISP2 (p<0.001).

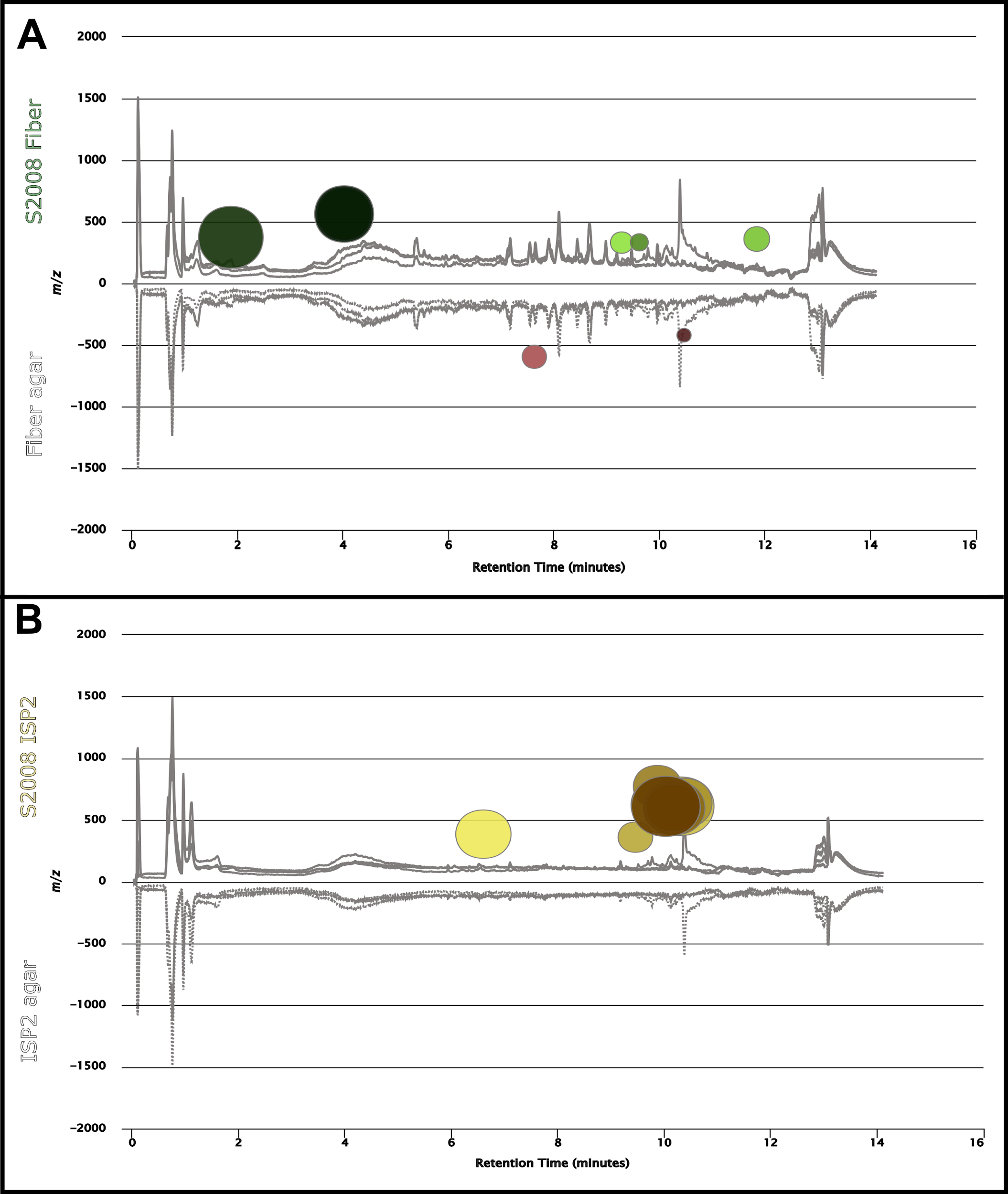

Table S1: Library isolates and quality of data collection. This table shows all of the library isolates and whether data was collected on the isolates and the quality of the data collected. If the signal to noise ratio was less than 15:1, data was considered poor. This table also highlights which isolates we were unable to revive in liquid culture or on solid agar after initial isolation from gut microbe cultivation.

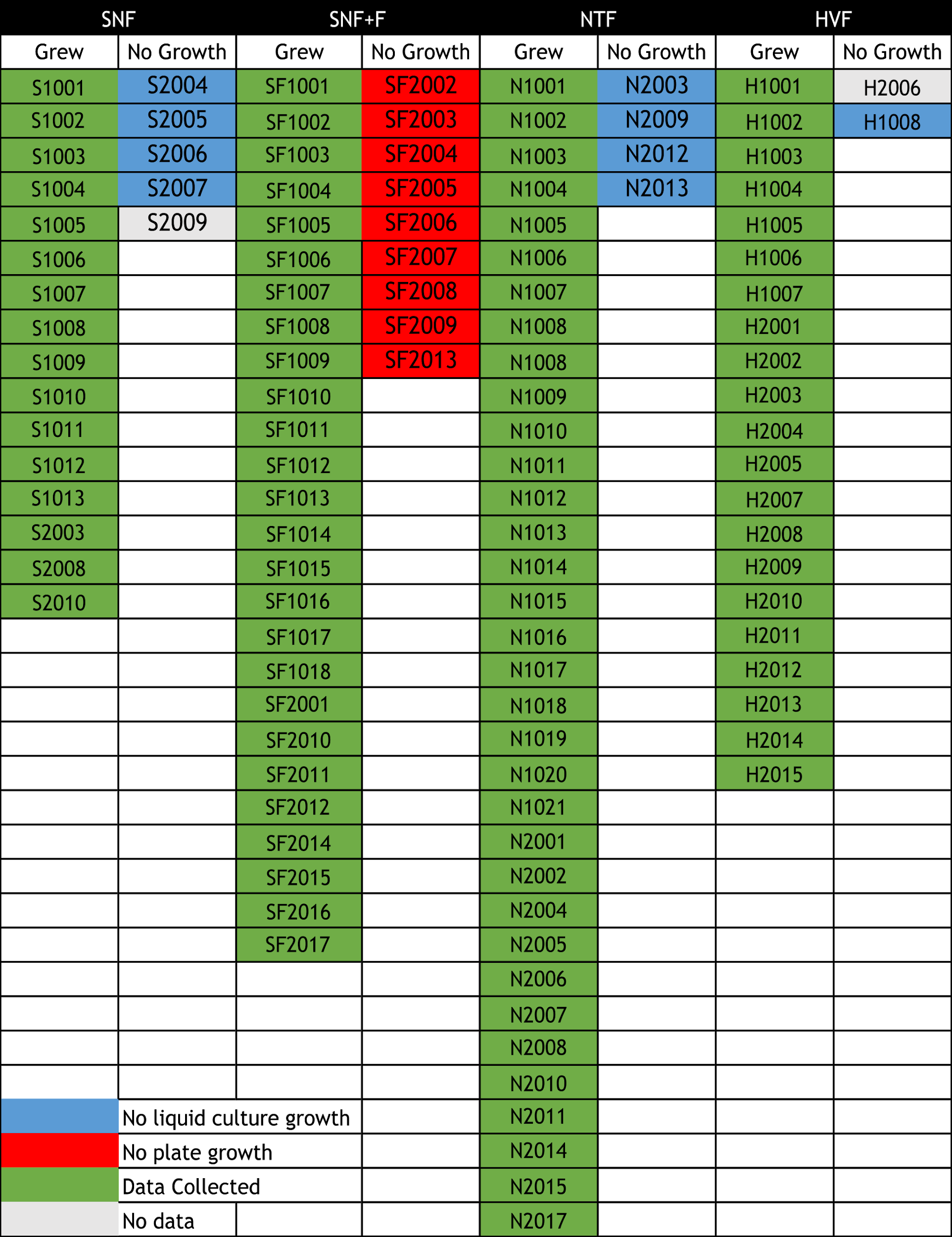

Table S2: Medium recipes

| **HVF**    1 L Milli-Q Water  20g agar  10g starch  1.7 KCl  0.01g FeS04*7H2O  0.3g KNO3  0.5g MgSO4*7H2O  0.5g Na2HPO4  (Vitamin Mixture from Federle Lab):  0.2mg p-aminobenzoic acid  0.2mg biotin  0.8mg folic acid  1mg niacinamide  2.5mg beta- nicotinamide adenine dinucleotide  2mg panthothenate calcium salt  1mg pyridoxal  1mg pyridoxamine dihydrochloride  2mg riboflavin  1mg thiamine hydrochloride  0.1mg vitamin B12 | **SNF**    1 L Milli-Q Water  18g agar |
| --- | --- |
|  | **SNF + Fiber**    1 L Milli-Q Water  18g agar  20g fiber |
|  | **NTF**    1 L Milli-Q Water  20g agar  20g starch  0.5g NaCl  0.01g FeS04*7H2O  0.5g MgSO4*7H2O  0.5g K2HPO4  1g KNO3 |

Table S3: Results from 16S sequencing analysis of 58 library isolate and respective accession numbers.

| **Isolate** | **Strain** | **Accession number** | **Isolation media** |
| --- | --- | --- | --- |
| N1008 | *Kocuria* sp. | MN588220 | NTF |
| H1006 | *Bacillus subtilis* | MN588221 | HVF |
| N1012 | *Bacillus subtilis* | MN588222 | NTF |
| N1013 | *Rhodococcus* sp. | MN588223 | NTF |
| N1014 | *Bacillus subtilis* | MN588224 | NTF |
| N1018 | *Bacillus subtilis* | MN588225 | NTF |
| N1019 | *Bacillus subtilis* | MN588226 | NTF |
| N1021 | *Bacillus subtilis* | MN588227 | NTF |
| N1015 | *Bacillus subtilis* | MN588228 | NTF |
| N1017 | *Bacillus subtilis* | MN588229 | NTF |
| S1011 | *Bacillus subtilis* | MN588230 | SNF |
| SF1011 | *Bacillus subtilis* | MN588231 | SNF + Fiber |
| N1006 | *Bacillus subtilis* | MN588232 | NTF |
| N1004 | *Bacillus subtilis* | MN588233 | NTF |
| N1002 | *Rhodococcus* sp. | MN588234 | NTF |
| SF2012 | *Bacillus subtilis* | MN588235 | SNF +Fiber |
| SF2001 | *Shewanella* sp. | MN588236 | SNF +Fiber |
| H2014 | *Aeromonas* sp. | MN588237 | HVF |
| H2013 | *Pseudomonas monteilii* | MN588238 | HVF |
| S1004 | *Bacillus subtilis* | MN588239 | SNF |
| SF1016 | *Bacillus coagulans* | MN588240 | SNF +Fiber |
| N1009 | *Bacillus subtilis* | MN588241 | NTF |
| N1010 | *Bacillus subtilis* | MN588242 | NTF |
| S2003 | *Bosea* sp. | MN588243 | SNF |
| N2002 | *Aeromonas* sp. | MN588244 | NTF |
| N2004 | *Staphylococcus hominis* | MN588245 | NTF |
| N2005 | *Pseudomonas monteilii* | MN588246 | NTF |
| N2006 | *Pseudomonas monteilii* | MN588247 | NTF |
| N2008 | *Pseudomonas monteilii* | MN588248 | NTF |
| N2011 | *Shewanella* sp. | MN588249 | NTF |
| H2001 | *Ensifer* sp. | MN588250 | HVF |
| H2002 | *Ensifer* sp. | MN588251 | HVF |
| H2003 | *Bosea* sp. | MN588252 | HVF |
| H2005 | *Mycolicibacterium* sp. | MN588253 | HVF |
| H2008 | *Bacillus licheniformis* | MN588254 | HVF |
| H2009 | *Ensifer* sp. | MN588255 | HVF |
| H2010 | *Ensifer* sp. | MN588256 | HVF |
| H2011 | *Bacillus flexus* | MN588257 | HVF |
| H2012 | *Bacillus flexus* | MN588258 | HVF |
| H1001 | *Bacillus subtilis* | MN588259 | HVF |
| H1004 | *Bacillus* sp. | MN588260 | HVF |
| S1008 | *Bacillus* sp. | MN588261 | SNF |
| S1013 | *Bacillus subtilis* | MN588262 | SNF |
| SF1003 | *Rhodococcus* sp. | MN588263 | SNF +Fiber |
| SF1005 | *Bacillus coagulans* | MN588264 | SNF +Fiber |
| SF1007 | *Bacillus subtilis* | MN588265 | SNF +Fiber |
| SF1008 | *Bacillus subtilis* | MN588266 | SNF +Fiber |
| SF1009 | *Bacillus subtilis* | MN588267 | SNF +Fiber |
| SF1012 | *Bacillus subtilis* | MN588268 | SNF +Fiber |
| SF1013 | *Bacillus subtilis* | MN588269 | SNF +Fiber |
| SF2017 | *Bacillus subtilis* | MN588270 | SNF +Fiber |
| SF1018 | *Bacillus coagulans* | MN588271 | SNF +Fiber |
| S2002 | *Bacillus licheniformis* | MN588272 | SNF |
| SF1015 | *Bacillus coagulans* | MN588273 | SNF +Fiber |
| S2001 | *Staphylococcus capitis* | MN588274 | SNF |
| SF1002 | *Bacillus coagulans* | MN588275 | SNF +Fiber |
| SF1014 | *Bacillus coagulans* | MN588276 | SNF +Fiber |

**Data Availability**

Massive Accession # for Raw MALDI data: MSV000084061

Massive Accession # for IDBac Files: MSV000084064
